# Assessing Confidence in the Results of Network Meta-Analysis (Cinema)

**DOI:** 10.1101/597047

**Authors:** Adriani Nikolakopoulou, Julian PT Higgins, Theodore Papakonstantinou, Anna Chaimani, Cinzia Del Giovane, Matthias Egger, Georgia Salanti

## Abstract

Evaluation of the credibility of results from a meta-analysis has become an intrinsic part of the evidence synthesis process. We present a methodological framework to evaluate Confidence In the results from Network Meta-Analysis (CINeMA) when multiple interventions are compared. CINeMA considers six domains and we outline the methods used to form judgements about within-study bias, across-studies bias, indirectness, imprecision, heterogeneity and incoherence. Key to judgements about within-study bias and indirectness is the percentage contribution matrix, which shows how much information each study contributes to the results from network meta-analysis. The use of contribution matrix allows the semi-automation of the process, implemented in a freely available web application (cinema.ispm.ch). In evaluating imprecision, heterogeneity and inconsistency we consider the impact of these components of variability in forming clinical decisions. Via three examples, we show that CINeMA improves transparency and avoids the selective use of evidence when forming judgements, thus limiting subjectivity in the process. CINeMA is easy to apply even in large and complicated networks, like a network involving 18 different antidepressant drugs.

## Introduction

Network meta-analysis has become an increasingly popular tool for developing treatment guidelines and making recommendations on reimbursement. However, less than one per cent of published network meta-analyses assess the credibility of their conclusions (1). The Grading of Recommendations Assessment, Development and Evaluation (GRADE) approach requires such an assessment of the confidence in the results from systematic reviews and meta-analyses, and many organizations, including the World Health Organization (WHO), have adopted the GRADE approach (2,3). Based on GRADE, two systems have been proposed to evaluate the credibility of results from network meta-analyses (4,5). However, the complexity of the methods and lack of suitable software have limited their uptake.

In this article we introduce the methodology underpinning the CINeMA approach (Confidence In Network Meta-Analysis), and present the advances that have recently been implemented in a freely available web application (*cinema.ispm.ch*) (6). CINeMA is based on the GRADE framework, with several conceptual and semantic differences (5). It covers six confidence domains: within-study bias (referring to the impact of risk of bias in the included studies), across-studies bias (referring to publication and other reporting bias), indirectness, imprecision, heterogeneity and incoherence. CINeMA assigns judgements at three levels (no concerns, some concerns or major concerns) to each of the six domains. Judgements across the six domains are then summarized to obtain four levels of confidence for each relative treatment effect, corresponding to the usual GRADE approach: very low, low, moderate or high.

Most network meta-analyses include only randomized controlled trials (RCTs), so we will focus on this study design, and on relative treatment effects. A network meta-analysis involves the integration of direct and indirect evidence in a network of relevant trials. We assume that evaluation of the credibility of results takes place once all primary analyses and sensitivity analyses have been undertaken. We assume that reviewers have implemented their pre-specified study inclusion criteria, which may include risk of bias considerations, and have obtained the best possible estimates of relative treatment effects using appropriate statistical methods (e.g. those described in (7–10)). The question is then how to make judgements about the credibility of relative treatment effects, given that trials with variable risk of bias, precision, relevance and heterogeneity contribute information to the estimate.

This paper addresses how judgements should be formed about the six CINeMA domains. We illustrate the methods using three examples: a network of trials that compare outcomes of various diagnostic strategies in patients with suspected acute coronary syndrome (11), a network of trials comparing the effectiveness of 18 antidepressants for major depression (12), and a network comparing adverse events of statins (13). The three examples are introduced in <u>**Error! Reference source not found**</u>.. All analyses were done in R software using the *netmeta* package and the CINeMA web application (Box 2) (6,14).

## Within-Study Bias

### Background and Definitions

Within-study bias refers to shortcomings in the design or conduct of a study that can lead to an estimated relative treatment effect that systematically differs from the truth. In our framework we assume that studies have been assessed for risk of bias. The majority of published systematic reviews of RCTs currently use a tool developed by Cochrane to evaluate risk of bias (15). This tool classifies studies as having low, unclear or high risk of bias for various bias components (such as allocation concealment, attrition, blinding etc.), and these judgements are then summarized across domains. A revision of the tool takes a similar approach but labels the levels as low risk of bias, some concerns and high risk of bias (16).

### The Cinema Approach

While it is straightforward to gauge the impact of within-study biases on the summary relative treatment effect in a pairwise meta-analysis (17), in network meta-analysis studies contribute data to the estimation of each summary effect in a complex manner. In the first example discussed below we show the complexity underpinning the flow of information in the network of diagnostic modalities used to detect coronary artery disease. A treatment comparison directly evaluated in studies with low risk of bias might also be estimated indirectly (via a common comparator) using studies at high risk of bias, and vice versa. While studies at low risk of bias are expected to provide more credible results, it is often impractical to restrict the analysis to such studies. The treatment comparison of interest might not have been tested directly in any trial, or tested in only a few small trials with high risk of bias. Thus, even when direct evidence is present, judgements about the relative treatment effect cannot ignore the risk of bias in the studies providing indirect evidence.

If direct evidence is supplemented by indirect evidence via exactly one intermediate comparator, the risk of bias in such a one-step loop is considered along with the direct evidence. In complex networks, indirect evidence is often obtained via several routes, including one-step loops and loops involving several steps (see example). In general, it is not desirable to derive judgements by considering only the risk of bias in studies in a single one-step loop (4,18). This is because most studies in a network contribute *some* indirect information to every estimate of a relative treatment effect. Studies contribute more when their results are precise (e.g. large studies), when they provide direct evidence or when the indirect evidence does not involve many “steps”. For example, studies in a one-step indirect comparison contribute more than studies of the same precision in a two-step indirect comparison. We can quantify the contribution made by each study to each relative treatment effect on a 0 to 100 percent scale. These quantities can be written as a ‘percentage contribution matrix’, as shown elsewhere (19).

CINeMA combines the studies’ contributions with the risk of bias judgements to evaluate study limitation for each estimate of a relative treatment effect from a network meta-analysis. It uses the percentage contribution matrix to approximate the contribution of each study and then stratifies the percentage contribution from studies judged to be at low, moderate and high risk of bias. Using different colors, study limitations in direct comparisons can be shown graphically in the network plot, while study limitations in the estimates from a network meta-analysis are presented for each comparison in bar charts.

### Example: Comparing diagnostic modalities to detect coronary artery disease

Consider the comparison of Exercise ECG versus Standard care (Box 1). The direct evidence from a single study is at low risk of bias (3-arm study 12); so there are no study limitations when interpreting the direct odds ratio of 0.42 (Table 1). However, the odds ratio 0.52 from the network meta-analysis is estimated also by using indirect information via seven studies that compare standard care and CCTA and one study comparing exercise ECG and CCTA. Additionally, we have indirect evidence via stress echo. The risk of bias in these eleven studies providing indirect evidence varies. Every study in the two one-step loops contributes information proportional to its precision (the inverse of the squared standard error, largely driven by sample size). Consequently, some judgement about study limitations for the indirect evidence can be made by considering that a there is a large amount of information from studies at high risk of bias (2162 participants randomized) and low risk of bias (2788 participants) and relatively little information from studies at moderate risk of bias (362 participants). Direct evidence from the small study number 12 (130 participants) at low risk of bias is considered separately, as it has greater influence than the indirect evidence.

#### Box 1.

Description of three network meta-analyses used to illustrate the CINeMA approach to assess confidence in network meta-analysis.

##### Diagnostic strategies for patients with low risk of acute coronary syndrome

Siontis et al performed a network meta-analysis to of randomized trials to evaluate the differences between the non-invasive diagnostic modalities used to detect coronary artery disease in patients presenting with symptoms suggestive of acute coronary syndrome (11). Differences between the diagnostic modalities were evaluated with respect to the number of downstream referrals for invasive coronary angiography and other clinical outcomes. For outcome referrals, 18 studies were included. The network is presented in Figure 1A and the data in Table S1. The results from the network meta-analysis are presented in Table 1.

##### Antidepressants for moderate and major depression

Cipriani et al compared 18 commonly prescribed antidepressants, which were studied in 179 head-to-head randomized trials involving patients diagnosed with major/moderate depression (12). The primary efficacy outcome was response measured as 50% reduction in the symptoms scale between baseline and 8 weeks of follow-up. According to the inclusion criteria specified in the protocol only studies at low or moderate risk of bias were included (58). The methodological and statistical details presented in the published article and its appendix. Here, we will focus on how judgements about credibility of the results were derived. The network is presented in Figure 1B and the data is available in Mendeley Data (DOI:10.17632/83rthbp8ys.2).

##### Comparative tolerability and harms of statins

The aim of the systematic review by Naci et al. (37) was to determine the comparative tolerability and harms of eight statins. The outcome considered here is the number of patients who discontinued therapy due to adverse effects, measured as an odds ratio. This outcome was evaluated in 101 studies. The network is presented in Figure 1C and the outcome data are given in Table S4. The results of the network meta-analysis are presented in Table S5 and the results from SIDE splitting in Table S6.

**Table 1.** Results from pairwise (upper triangle) and network meta-analysis (lower triangle) from the network of non-invasive diagnostic strategies for the detection of coronary artery disease in Figure 1A. Odds ratios and their 95% confidence intervals are presented for referrals for invasive coronary angiography. Odds ratios in the lower triangle less than one favor the strategy in the column; odds ratios in the upper triangle less than one favor the strategy in the row. Cells with a dot indicate that no direct studies examine the particular comparison.

|  |  |  |  |  |  |
| --- | --- | --- | --- | --- | --- |
| **CCTA** | . | 2.25<br>[1.04 - 4.90] | 1.04<br>[0.70 - 1.55] | 1.23<br>[1.00 - 1.50] | . |
| 3.07<br>[1.46 - 6.45] | **CMR** | . | . | 0.38<br>[0.18 - 0.78] | . |
| 2.24<br>[1.22 - 4.11] | 0.73<br>[0.28 - 1.88] | **Exercise ECG** | . | 0.42<br>[0.14 - 1.30] | 1.93<br>[1.39 - 2.67] |
| 1.27<br>[1.01 - 1.60] | 0.42<br>[0.20 - 0.87] | 0.57<br>[0.30 - 1.07] | **SPECT-MPI** | 0.87<br>[0.71 - 1.06] | . |
| 1.17<br>[0.97 - 1.40] | 0.38<br>[0.18 - 0.78] | 0.52<br>[0.28 - 0.96] | 0.92<br>[0.76 - 1.10] | **Standard Care** | 2.95<br>[0.97 - 8.98] |
| 4.31<br>[2.23 - 8.32] | 1.40<br>[0.53 - 3.74] | 1.93<br>[1.39 - 2.66] | 3.38<br>[1.71 - 6.68] | 3.69<br>[1.90 - 7.17] | **Stress Echo** |

*ECG: electrocardiogram; echo: echocardiography; SPECT-MPI: single photon emission computed* *tomography-myocardial perfusion imaging; CCTA: coronary computed tomographic angiography;* *CMR: cardiovascular magnetic resonance.*

Calculations become more complicated because studies in the indirect comparisons contribute information not only proportional to their study precision but also to their location in the network. Indirect evidence about exercise ECG versus SPECT-MPI comes from two one-step loops (via CCTA or via Standard Care) and three two-step loops (via CCTA-Standard Care, Stress Echo-Standard Care, Standard Care-CCTA) (Figure 1A). In each loop of evidence, a different subgroup of studies contributes indirect information and their sizes and risks of bias vary. For the odds ratio from the network meta-analysis comparing exercise ECG and SPECT-MPI, study 2 with sample size 400 will be more influential than study 8 (with sample size 1392) because study 2 contributes one-step indirect evidence (via standard care).

**Figure 1.**
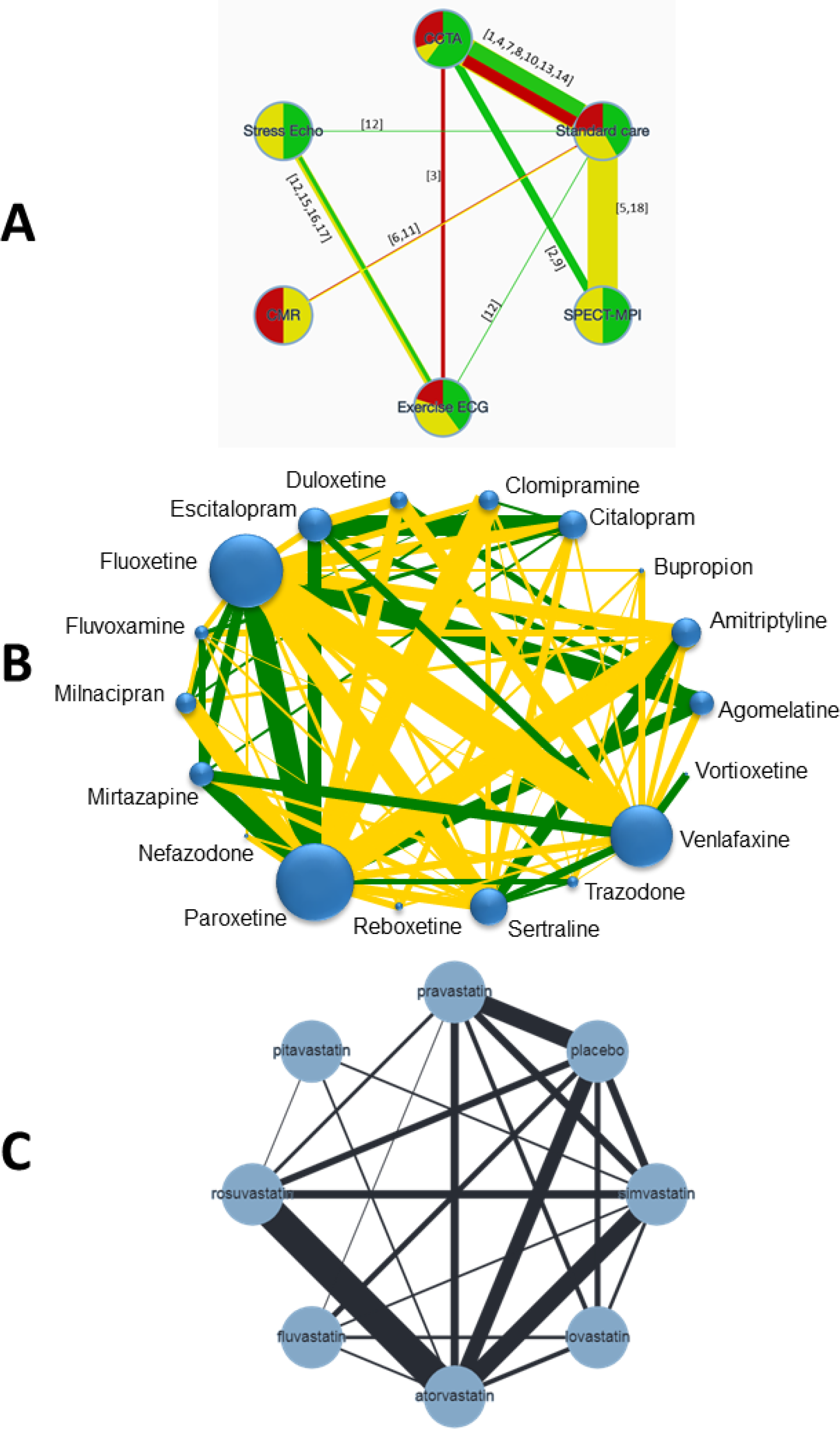
Network plots of the three network meta-analyses used as examples. The width of the edges are proportional to the number of patients randomised in each comparison. A: Network of randomised controlled trials comparing non-invasive diagnostic strategies for the detection of coronary artery disease in patients with low risk acute coronary syndrome. The colours of edges and nodes refer to the risk of bias; low (green), moderate (yellow) and red (high). In square brackets are the study IDs as presented in Table S1. B: Network of randomised controlled trials comparing active antidepressants in patients with moderate/major depression. The colours of edges refer to the risk of bias; low (green), moderate (yellow) and red (high). The size of nodes is proportional to the number of studies examining each treatment. C: Network of randomised controlled trials comparing statins with respect to adverse effects. ECG: electrocardiogram; echo: echocardiography; SPECT-MPI: single photon emission computed tomography-myocardial perfusion imaging; CCTA: coronary computed tomographic angiography; CMR: cardiovascular magnetic resonance.

Table 2 shows the percentage contribution matrix for the network and the columns represent the studies, grouped by comparison. The rows represent all relative treatment effects from network meta-analysis. The matrix entries show how much each study contributes to the estimation of each relative treatment effect. This information combined with the risk of bias judgements can be presented as a bar chart, as shown in Figure 2. Now, it is much easier to judge study limitations for each odds ratio; the larger the contribution from studies at high or moderate risk of bias, the more concerned we are about study limitations. Using this graph, we can infer that the total evidence from the network meta-analysis for the comparison of exercise ECG with SPECT-MPI involves low, moderate and high risk of bias studies with percentages 44%, 32% and 24%, respectively.

**Table 2.** The percentage contribution matrix for the network presented in Figure 1A. The columns refer to the studies (grouped by comparison) and the rows refer to the relative treatment effects (grouped into mixed and indirect evidence) from network meta-analysis. The entries show how much each study contributes (as percentage) to the estimation of relative treatment effects.

| Direct comparisons (number of studies) | CCTA vs Exercise ECG (1) | CCTA vs SPECT-MPI (2) |  | CCTA vs Standard care (7) |  |  |  |  |  |  | CMR vs Standard care (2) |  | Exercise ECG vs Standard care (1) | Exercise ECG vs Stress Echo (4) |  |  |  | SPECT-MPI vs Standard care (2) |  | Standard care vs Stress Echo (1) |
| --- | --- | --- | --- | --- | --- | --- | --- | --- | --- | --- | --- | --- | --- | --- | --- | --- | --- | --- | --- | --- |
| NMA Estimates/study IDs | 3 | 2 | 9 | 1 | 10 | 13 | 14 | 4 | 7 | 8 | 11 | 6 | 12 | 12 | 15 | 16 | 17 | 18 | 5 | 12 |
| Mixed estimates |  |  |  |  |  |  |  |  |  |  |  |  |  |  |  |  |  |  |  |  |
| CCTA:Exercise ECG | 52 | 1 | 1 | 3 | 0 | 3 | 1 | 3 | 4 | 4 | 0 | 0 | 14 | 0 | 3 | 0 | 2 | 1 | 1 | 6 |
| CCTA:SPECT-MPI | 1 | 18 | 16 | 5 | 1 | 5 | 1 | 6 | 7 | 7 | 0 | 0 | 0 | 0 | 0 | 0 | 0 | 22 | 10 | 0 |
| CCTA:Standard care | 1 | 4 | 4 | 13 | 2 | 13 | 3 | 15 | 18 | 17 | 0 | 0 | 1 | 0 | 0 | 0 | 0 | 6 | 3 | 0 |
| CMR:Standard care | 0 | 0 | 0 | 0 | 0 | 0 | 0 | 0 | 0 | 0 | 60 | 40 | 0 | 0 | 0 | 0 | 0 | 0 | 0 | 0 |
| Exercise ECG:Standard care | 23 | 1 | 1 | 3 | 0 | 3 | 1 | 4 | 5 | 4 | 0 | 0 | 30 | 1 | 6 | 1 | 3 | 2 | 1 | 11 |
| Exercise ECG:Stress Echo | 1 | 0 | 0 | 0 | 0 | 0 | 0 | 0 | 0 | 0 | 0 | 0 | 1 | 5 | 52 | 8 | 29 | 0 | 0 | 2 |
| SPECT-MPI:Standard care | 0 | 5 | 4 | 1 | 0 | 1 | 0 | 2 | 2 | 2 | 0 | 0 | 0 | 0 | 0 | 0 | 0 | 57 | 26 | 0 |
| Standard care:Stress Echo | 14 | 1 | 1 | 2 | 0 | 2 | 1 | 2 | 3 | 3 | 0 | 0 | 14 | 2 | 16 | 2 | 9 | 1 | 1 | 27 |
| Indirect estimates | -- | -- | -- | -- | -- | -- | -- | -- | -- | -- | -- | -- | 0 | 0 | -- | -- | -- | -- | -- | 0 |
| CCTA:CMR | 1 | 3 | 2 | 6 | 1 | 7 | 2 | 8 | 9 | 8 | 28 | 19 | 1 | 0 | 0 | 0 | 0 | 4 | 2 | 0 |
| CCTA:Stress Echo | 24 | 1 | 1 | 3 | 0 | 3 | 1 | 3 | 4 | 4 | 0 | 0 | 8 | 2 | 18 | 3 | 10 | 1 | 1 | 13 |
| CMR:Exercise ECG | 16 | 1 | 1 | 2 | 0 | 2 | 1 | 3 | 3 | 3 | 22 | 15 | 15 | 0 | 4 | 1 | 2 | 1 | 1 | 7 |
| CMR:SPECT-MPI | 0 | 3 | 3 | 1 | 0 | 1 | 0 | 1 | 1 | 1 | 28 | 19 | 0 | 0 | 0 | 0 | 0 | 28 | 13 | 0 |
| CMR:Stress Echo | 11 | 1 | 1 | 1 | 0 | 2 | 0 | 2 | 2 | 2 | 20 | 14 | 9 | 1 | 11 | 2 | 6 | 1 | 0 | 13 |
| Exercise ECG:SPECT-MPI | 21 | 7 | 6 | 1 | 0 | 1 | 0 | 1 | 2 | 2 | 0 | 0 | 15 | 0 | 4 | 1 | 2 | 20 | 9 | 7 |
| SPECT-MPI:Stress Echo | 14 | 5 | 4 | 1 | 0 | 1 | 0 | 1 | 1 | 1 | 0 | 0 | 9 | 1 | 13 | 2 | 7 | 19 | 9 | 13 |

ECG: electrocardiogram; echo: echocardiography; SPECT-MPI: single photon emission computed tomography-myocardial perfusion imaging; CCTA: coronary computed tomographic angiography; CMR: cardiovascular magnetic resonance.

**Figure 2.**
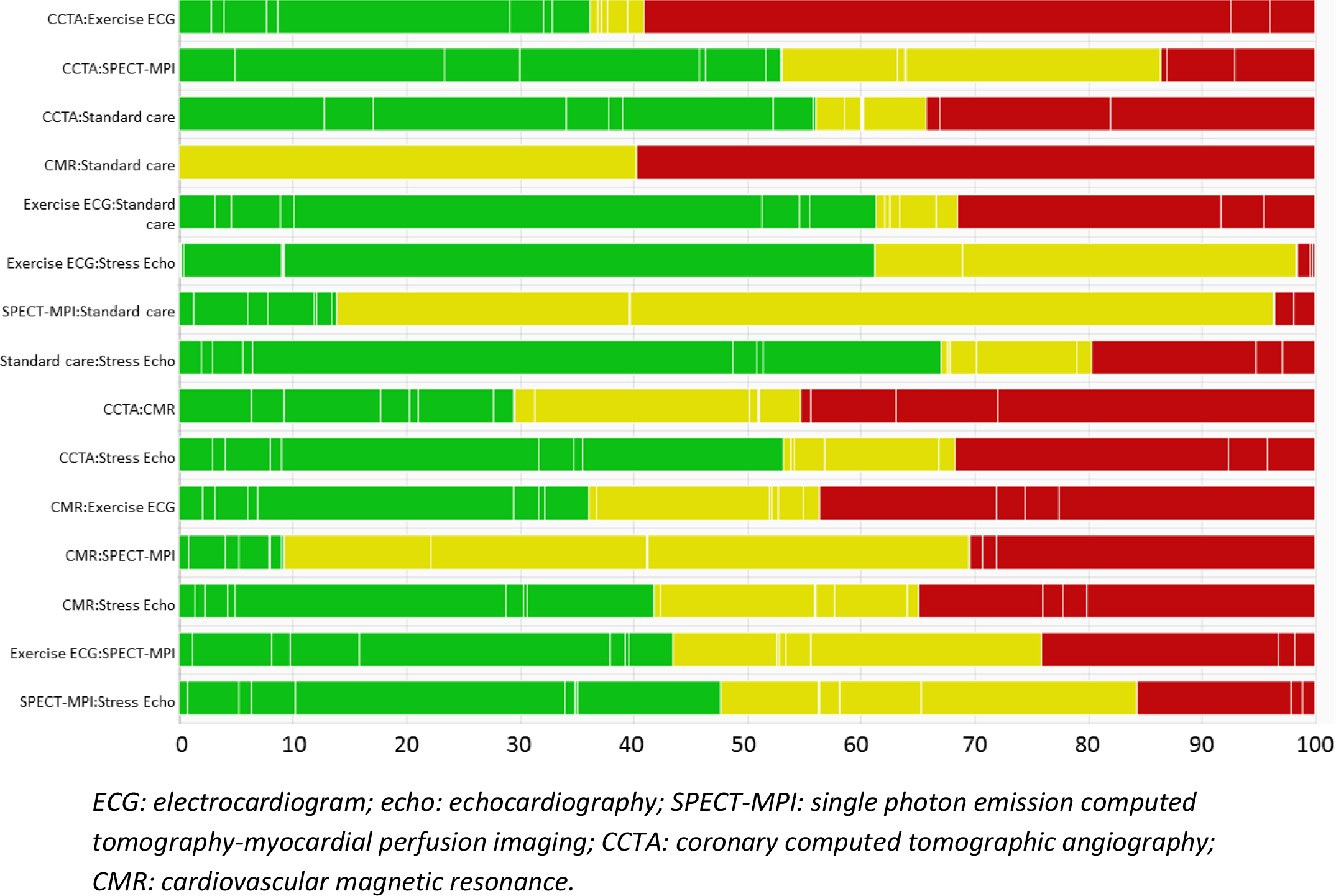
Risk of bias bar chart for the comparison of non-invasive diagnostic strategies for the detection of coronary artery disease. Each bar represents a relative treatment effect estimated using the data in the network in Error! Reference source not found.A. White vertical lines indicate the percentage contribution of separate studies. Each bar shows the percentage contribution from studies judged to be at low (green), moderate (yellow) and high (red) risk of bias.

The CINeMA software offers the option to automate production of judgments, based on the data presented in these bar graphs combined with specific rules. One possible rule is to compute a weighted average level of risk of bias, assigning scores of −1, 0 and 1 to low, moderate and high risk of bias. For the comparison exercise ECG vs SPECT-MPI, this would produce a weighted score of 0.44 × (−1) + 0.32 × 0 + 0.24 × 1 = −0.20, which corresponds to some concerns in the scoring scheme.

### Example: comparing antidepressants

We will focus on evaluating the results for three comparisons; amitriptyline vs milnacipran (one direct study at low and one at moderate risk of bias), mirtazapine versus paroxetine (three direct studies at low risk of bias and two at moderate) and amitriptyline vs clomipramine (no direct studies). The odds ratios for treatment response are presented in Table 3. We use this example to illustrate the use of sensitivity analysis and how it can inform the amount of contribution of studies at moderate and high risk of bias that we can tolerate.

**Table 3.** Summary odds ratios from network meta-analysis comparing six antidepressants and sensitivity analyses excluding studies at moderate risk of bias.

| Comparison | Response odds ratio [95% confidence interval] |  |
| --- | --- | --- |
|  | *All studies*<br>*(179 studies)* | *Studies at low risk of bias*<br>*(83 studies)* |
| Amitriptyline versus Milnacipran | 1.11 [0.85; 1.43] | 1.10 [0.77; 1.59] |
| Mirtazapine versus Paroxetine | 1.07 [0.88; 1.30] | 1.08 [0.83; 1.39] |
| Amitriptyline versus Clomipramine | 1.24 [0.97; 1.59] | 0.96 [0.59; 1.57] |

For the first two treatment comparisons in Table 3, the contribution from studies at low risk of bias is more than 50%. Moreover, the sensitivity analysis excluding studies at moderate risk of bias provides results comparable to those obtained from all studies. Thus, one can derive the judgment of no concerns for amitriptyline versus milnacipran and mirtazapine versus paroxetine. However, the estimation of the relative treatment effect of amitriptyline versus clomipramine comes by more than 60% from studies at moderate risk of bias. Given also that the odds ratio from the sensitivity analysis is quite different from to the one obtained from all studies, we judge as some concerns the amitriptyline versus clomipramine comparison.

## Across-studies bias

### Background and definitions

Across-studies bias occurs when the studies included in the systematic review are not a representative sample of the studies undertaken. This phenomenon can be the result of the suppression of statistically significant (or “negative”) findings (publication bias), their delayed publication (time-lag bias) or omission of unfavorable study results (outcome reporting bias). The presence and the impact of such biases has been well documented (20–26). Across-studies bias is a missing data problem, and hence it is impossible to conclude with certainty for or against its presence in a given dataset. Consequently, and in agreement with the GRADE system, CINeMA assumes two possible descriptions for across-studies bias: suspected and undetected.

### The CINeMA approach

Assessment of the risk of across-studies bias follows considerations on pairwise meta-analysis (27). Conditions associated with ‘suspected’ across-studies bias include:

- Failure to include unpublished data and data from grey literature.
- The meta-analysis is based on a small number of positive early findings, for example for a drug newly introduced on the market (as early evidence is likely to overestimate its efficacy and safety) (27).
- The treatment comparison is studied exclusively or primarily in industry-funded trials (28,29).
- There is previous evidence documenting the presence of reporting bias. For example the study by Turner et al. documented publication bias in placebo-controlled antidepressant trials (30).

Across-studies bias is considered ‘undetected’ when

- Data from unpublished studies have been identified and their findings agree with those in published studies
- There is a tradition of prospective trial registration in the field and protocols or clinical trial registries do not indicate important discrepancies with published reports
- Empirical examination of patterns of results between small and large studies, using the comparison-adjusted funnel plot (31,32), regression models (33) or selection models (34) do not indicate that results from small studies differ from those in published studies.

### Example: comparing antidepressants

The literature search retrieved supplementary and unpublished information from clinical trial registries, regulatory agencies’ repositories and drug companies’ websites (particularly for the newest and most recently marketed antidepressants). Results from published and unpublished studies did not differ materially, no asymmetry was observed in the funnel plot (12) and meta-regression did not indicate an association between study precision and study odds ratio. However, the authors decided that they cannot completely rule out the possibility that some studies are missing because the field of antidepressant trials has been shown to be prone to publication bias. Consequently, the review team decided to assume that across-studies bias was ‘suspected’ for all drug comparisons.

## Indirectness

### Background and definitions

Systematic reviews are based on a focused research question, with a clearly defined population, intervention and setting of interest. In the GRADE framework for pairwise meta-analysis, indirectness refers to the relevance of the included studies to the research question (35). Study populations, interventions, outcomes and study settings should match the inclusion criteria of the systematic review but might not be representative of the settings, populations or outcomes about which reviewers want to make inferences. For example, a systematic review aiming to provide evidence about treating middle-aged adults might identify studies in elderly patients; these studies will have an indirect relevance.

### The CINeMA approach

We suggest that each study included in the network is evaluated according to its relevance to the research question and classified into low, moderate or high indirectness. Note that only participant, intervention and outcome characteristics that are likely associated with the relative effect of an intervention against another (that is, effect modifying variables) should be considered. Then, the study-level judgments can be combined with the percentage contribution matrix to produce a bar plot similar to the one presented in Figure 2. Evaluation of indirectness for each relative treatment effect can then proceed by judging whether the contribution from studies of high or moderate indirectness is important.

This approach also addresses the assumption of transitivity in network meta-analysis. Transitivity assumes that we can learn about the relative treatment effect of, say treatment A versus treatment B from an indirect comparison via C. This holds when the distributions of all effect modifiers are comparable in A versus C and B versus C studies. Differences in the distribution of effect modifiers across studies and comparisons will indicate intransitivity. Evaluation of the distribution of effect modifiers is only possible when enough studies are available per comparison. Consequently, the proposed approach will not address intransitivity in sparse networks (when there are few studies compared to the total number of treatments). Assessment of transitivity will be challenging or impossible for interventions that are poorly connected to the network. A further potential obstacle is that details of important effect modifiers might not always be reported in trial reports. For these reasons, we recommend that the network structure and the amount of available data are considered, and that judgments are on the side of caution, as highlighted in the following example.

### Example: comparing antidepressants

Cipriani et al concluded that there is no indirectness in any of the studies included and that the distribution of modifiers was similar across studies and comparisons (12). However, they decided to downgrade evidence about drugs that are poorly connected to the network. For example, vortioxetine was examined in a single study and consequently it was difficult to assess the comparability of effect modifiers in the comparisons with vortioxetine. Consequently, Cipriani et al. voiced concerns about indirectness for all comparisons with vortioxetine.

## IMPRECISION

### Background and definitions

One of the key advantages of network meta-analysis compared to pairwise meta-analysis is the ability to gain precision (36): adding indirect evidence on a particular treatment comparison on top of direct evidence leads to narrower confidence intervals than using the direct evidence alone. However, in network meta-analysis treatment effects are also estimated with uncertainty, typically expressed as 95% confidence intervals that give an indication of where the true effect is likely to lie. To evaluate imprecision it is customary to define relative treatment effects that exclude any clinically important differences in outcomes between interventions (26). At its simplest, this treatment effect might correspond to no effect (0 on an additive scale, 1 on a ratio scale). This would mean that even a small difference is considered important, leading to one treatment being preferred over another. Alternatively, ranges may be defined that divide relative treatment effects into three categories: ‘in favour of A’, ‘no important difference between A and B’, and ‘in favour of B’. The middle range is the ‘range of equivalence’, which includes treatment effects that correspond to clinically unimportant differences between interventions. The range of equivalence can be symmetrical (when a clinically important difference is defined, and its reciprocal constitutes the clinically important difference in the opposite direction) or asymmetrical (when clinically important differences vary by direction of effect). For simplicity, we will assume symmetrical ranges of equivalence.

### The CINeMA approach

The approach to imprecision consists of comparing the range of treatment effects included in the 95% confidence interval with the range of equivalence. If the 95% confidence interval extends to differences in treatment effects that would lead to different conclusions, for example covering two or all three of the categories defined above, then the results would be considered imprecise, reducing confidence in the treatment effect estimate. Figure 3 shows a hypothetical forest plot that illustrates the CINeMA rules to assess imprecision of network treatment effect estimates for an odds ratio of 0.8. ‘Major concerns’ are assigned to NMA treatment effects with 95% confidence intervals that cross both limits of the range of equivalence, ‘some concerns’ if only the lower or the upper limit of the range of equivalence is crossed and ‘no concerns’ apply to estimates that do not cross either value.

**Figure 3.**
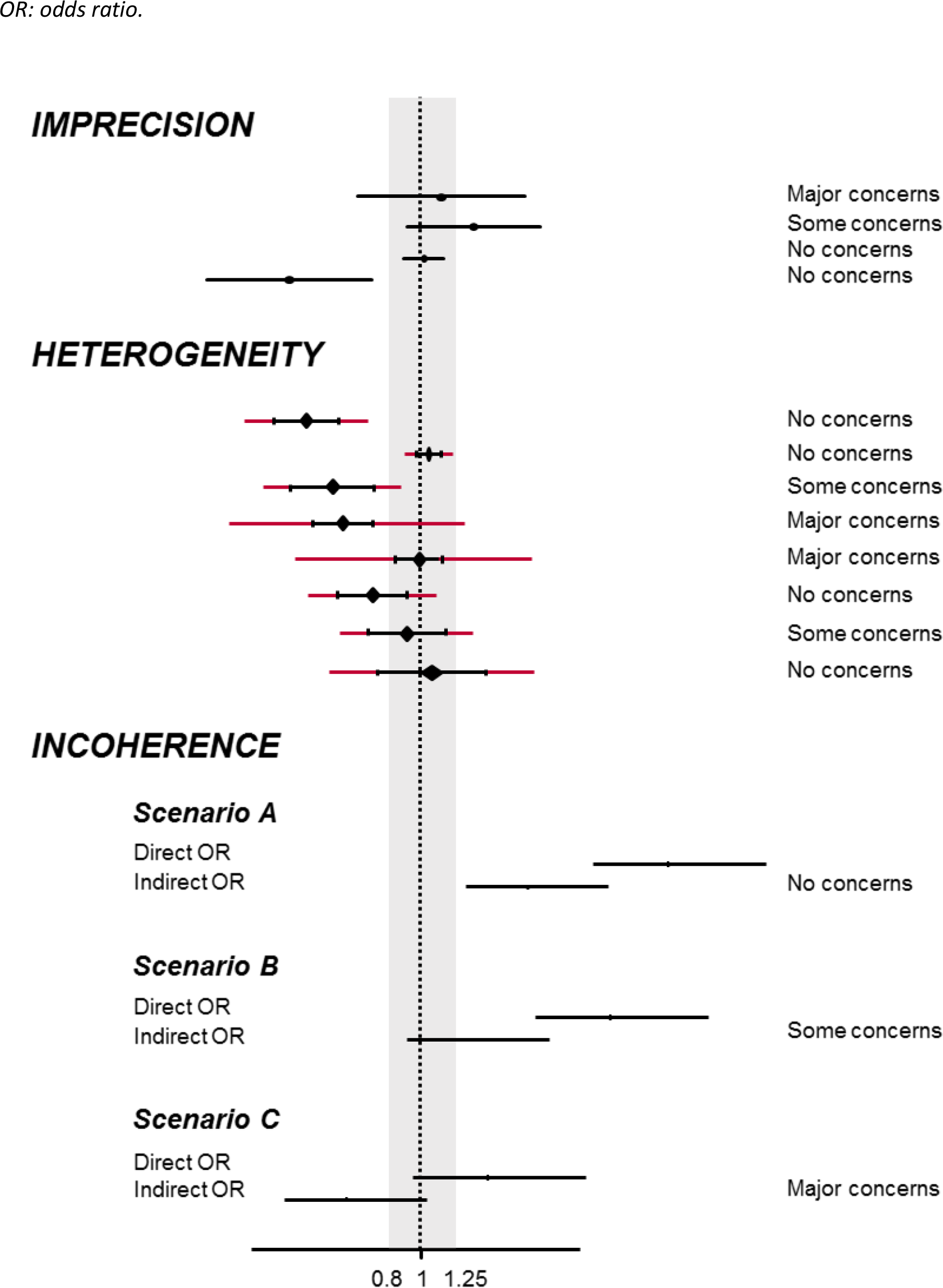
CINeMA rules to assess imprecision, heterogeneity and incoherence of network treatment effects. The range of equivalence is from 0.8 to 1.25. Black lines indicate confidence intervals and red lines indicate prediction intervals. For the three scenarios presented for incoherence, inconsistency factor is 1.27 (1.05 to 1.55).

### Example: adverse events of statins

Consider the network comparing adverse events of different statins, introduced in Box 1 and shown in Figure 1C (37). Let us assume a range of equivalence such that an odds ratio greater than 1.05 or below 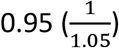 would lead to favouring one the two treatments. Odds ratios between 0.95 and 1.05 would be interpreted as no important differences in the safety profile of the two statins. The 95% confidence interval of pravastatin versus rosuvastatin is quite wide, including odds ratios from 1.09 to 1.82 (Figure 4), but any treatment effect in this range would lead to the conclusion that pravastatin is safer than rosuvastatin. Thus, in this case the imprecision does not reduce the confidence that can be placed in the comparison of pravastatin with rosuvastatin (‘no concerns’). The 95% confidence interval of pravastatin versus simvastatin is slightly wider (0.84 to 1.42) and, more importantly, the interval covers all three areas, i.e. favouring pravastatin, favouring simvastatin and no important difference. This result is very imprecise, and a rating of ‘major concerns’ applies. The comparison of rosuvastatin versus simvastatin is more certain, but it is again unclear which drug has fewer adverse effects: most estimates within the 95% confidence interval favour simvastatin, but the interval crosses into the range of equivalence. A rating of ‘some concerns’ will be appropriate here.

**Figure 4.**
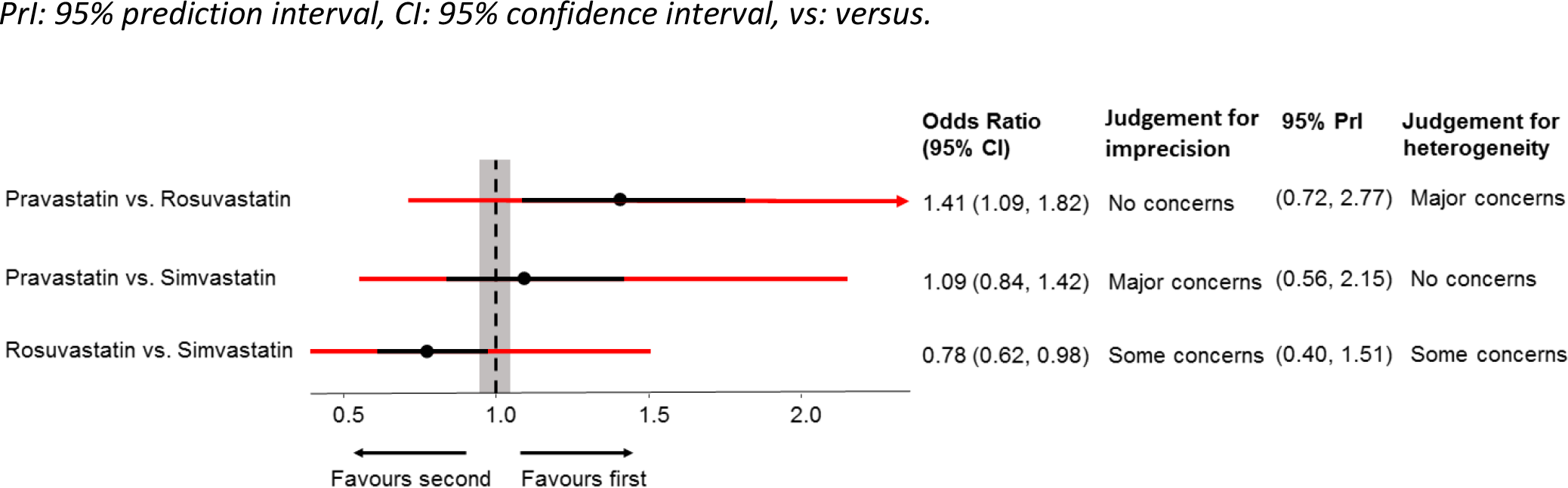
Network meta-analysis odds ratios from the network of statins their 95% confidence intervals (black lines) and their 95% prediction intervals (red lines). The range of equivalence is from 0.95 to 1.05.

### Example: efficacy of antidepressants

In the network of antidepressants, the authors defined clinically important effects as an odds ratio smaller than 0.8 and larger than its reciprocal 1.25 (12). We use this range of equivalence (0.8 to 1.25) in this example. We will concentrate on three comparisons, clomipramine versus fluvoxamine, citalopram versus venlafaxine and amitriptyline versus paroxetine (Table 4). The 95% confidence interval of the odds ratio comparing clomipramine with fluvoxamine (0.75 to 1.32) includes clinically important effects in both directions, implying large uncertainty in which drug should be favored (‘major concerns’) (Table 5). The odds ratio for citalopram versus venlafaxine is 1.12 (95% confidence interval 0.90 to 1.39), favoring venlafaxine, but the interval includes values within the range of equivalence. The verdict therefore is ‘some concerns’. Finally, the odds ratio of amitriptyline versus paroxetine is 0.96 (95% confidence interval 0.82 to 1.13) in favor of amitriptyline. Despite the fact that the estimate includes 1, it is not imprecise because the 95% confidence interval is within the range of equivalence (‘no concerns’).

**Table 4.**
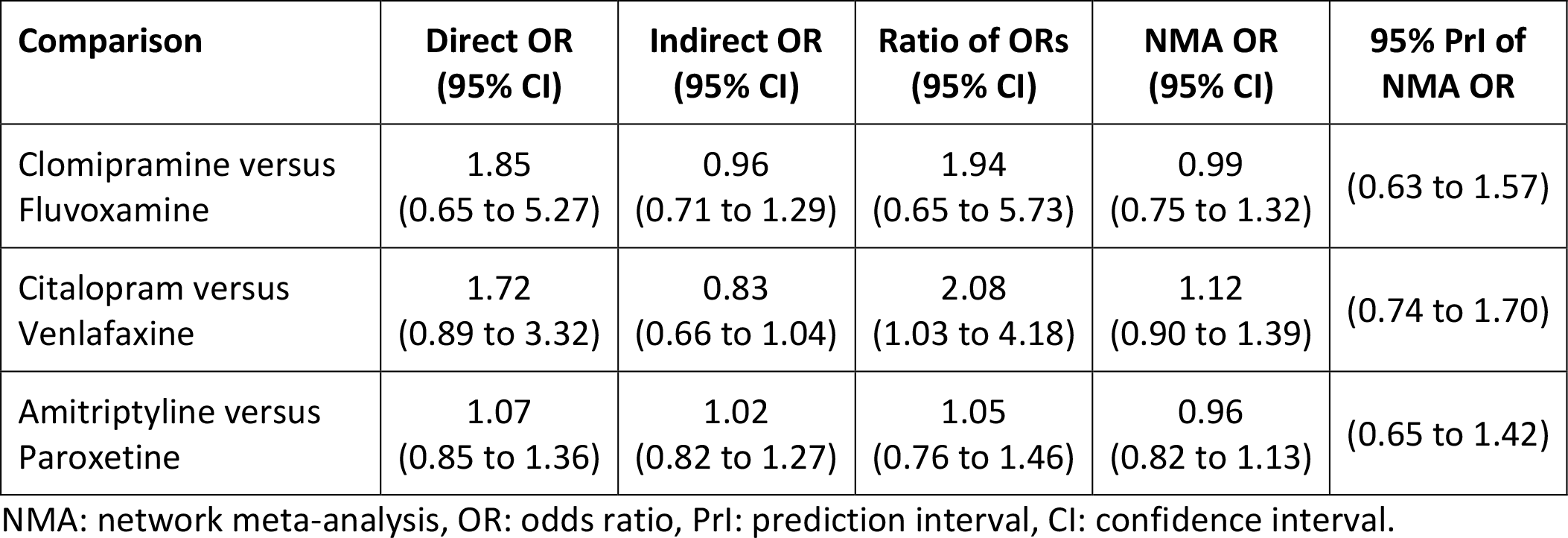
Results from direct, indirect and mixed evidence along with confidence and prediction intervals and incoherence ratio of odds ratios for the network of antidepressants. Odds ratios lower than 1 favour the first treatment.

**Table 5.** Level of concern for three network meta-analysis odds ratios from the network of antidepressants for the domains imprecision, heterogeneity and incoherence. See Table 4 for odds ratios.

| **Comparison** | **Imprecision** | **Heterogeneity** | **Incoherence** |
| --- | --- | --- | --- |
| Clomipramine versus Fluvoxamine | Major concerns | No concerns | No concerns |
| Citalopram versus Venlafaxine | Some concerns | Some concerns | Major concerns |
| Amitriptyline versus Paroxetine | No concerns | Major concerns | No concerns |

## Heterogeneity

### Background and definitions

Variability in the results of studies contributing to a particular comparison influences the confidence we have in the result for that comparison. If this variability reflects genuine differences between studies, rather than random variation, it is usually referred to as heterogeneity. The GRADE system for pairwise meta-analysis uses the term inconsistency to describe such variability (38). In network meta-analysis, there may be variation in the relative treatment effects between studies within a comparison, i.e. heterogeneity, or variation between direct and indirect sources of evidence across comparisons, i.e. incoherence (39–42) which we discuss in the next paragraph. The two notions are closely related; incoherence can be seen as a special form of heterogeneity.

There are several ways of measuring heterogeneity in a set of trials. The variance of the distribution of the underlying treatment effects (*τ*^2^), is a useful measure of the magnitude of heterogeneity. One can estimate heterogeneity variances from each pairwise meta-analysis and, under the usual assumption of a single variance across comparisons, a common heterogeneity variance for the whole network. The magnitude of *τ*^2^ is usefully expressed in a prediction interval, which shows where the true effect of a new study similar to the existing studies is expected to lie (28).

### The CINeMA approach

Similarly to imprecision, the CINeMA approach to heterogeneity considers its influence on clinical conclusions. Large variability in the included studies does not necessarily affect conclusions, while even small amounts of heterogeneity may be important in some cases. The concordance between assessments based on confidence intervals (which do not capture heterogeneity) and prediction intervals (which do capture heterogeneity) can be used to assess the importance of heterogeneity. For example, a prediction interval may include values that would lead to different conclusions than suggested by the CI; in such a case, heterogeneity would be considered having important implications. The hypothetical forest plot of Figure 3 serves as an illustration of the CINeMA rules to assess heterogeneity of treatment effects for a clinically important odds ratio of below 0.8 or above 1.25.

With only a handful of trials, one cannot adequately estimate the amount of heterogeneity: prediction intervals derived from meta-analyses with very few studies can be unreliable. In this situation it may be more reasonable to interpret an estimate of heterogeneity (and its uncertainty) using empirical distributions. Turner et al. and Rhodes et al. analyzed many meta-analyses of binary and continuous outcomes, categorized them according to the outcome and type of intervention and comparison, and derived empirical distributions of heterogeneity values (16, 17). These empirical distributions can help to interpret the magnitude of heterogeneity, complementary to considerations based on prediction intervals.

### Example: adverse events of statins

In the statins example (Figure 1C), we assumed that the range of equivalence was 0.95 to 1.05. The prediction interval of pravastatin versus simvastatin is wide (Figure 4). However, the confidence interval for this comparison already extended into clinically important effects in both directions; thus, the implications of heterogeneity is not important and does not change the conclusion. The confidence interval for pravastatin versus rosuvastatin lies entirely above the equivalence range and is consequently considered sufficiently precise. However, the corresponding prediction interval crosses both boundaries (0.95 and 1.05), and we therefore would have ‘major concerns’ about the impact of heterogeneity. Similar considerations result in ‘some concerns’ regarding heterogeneity for the comparison rosuvastatin versus simvastatin.

### Example: efficacy of antidepressants

In the antidepressants network, the estimated amount of heterogeneity is small (*τ*^2^ = 0.03). The prediction interval for clomipramine versus fluvoxamine does not add further uncertainty to clinical conclusions beyond that already represented by the confidence interval (Table 4), so we have ‘no concerns’ about heterogeneity for that comparison (Table 5). The prediction interval of citalopram versus venlafaxine extend into clinically important effects in both directions (0.74 to 1.70) while the confidence interval does not extend into values in favour of citalopram, thus suggesting potential implications of heterogeneity (‘some concerns’). We have ‘major concerns’ about the impact of heterogeneity for the comparison amitriptyline versus paroxetine, since the confidence interval lies entirely within the range of equivalence, whereas the prediction interval includes clinically important effects in favour of both treatments (0.65, 1.42).

## Incoherence

### Background and definitions

The assumption of transitivity stipulates that we can compare two treatments indirectly via an intermediate treatment node. Incoherence is the statistical manifestation of intransitivity; if transitivity holds, the direct and indirect evidence will be in agreement (45,46). Conversely, if estimates from direct and indirect evidence disagree we conclude that transitivity does not hold. There are two approaches to quantifying incoherence. The first comprises methods that examine the agreement between direct and indirect evidence for specific comparisons in the network, while the second includes methods that examine incoherence in the entire network. SIDE (Separate Indirect from Direct Evidence) or “node splitting” (39)) is an example of the first set of methods, which are often referred to as local methods. It compares direct and indirect evidence for each comparison and computes an inconsistency factor with a confidence interval. The inconsistency factor is calculated as the difference of the two estimates for an additive measure (e.g. log odds ratio, log risk ratio, standardized mean difference) or as the ratio of the two estimates for measures on the ratio scale. This method can be applied to comparisons that are informed by both direct and indirect evidence. Consider for example the hypothetical example in Figure 3 (Incoherence, Scenario A). The studies directly comparing the two treatments result in a direct odds ratio of 1.75 (1.5 to 2) while the rest studies of the network that provide indirect evidence to the particular comparison gives an indirect odds ratio of 1.37 (1.2 to 1.55). The disagreement between direct and indirect odds ratios is expressed as the ‘inconsistency factor’ (1.27) which can be used to construct a confidence interval (1.05 to 1.55) and a test statistic, here resulting to a p-value of 0.07. A simpler version of SIDE splitting considers a single loop in the network (loop-specific approach (47)). The second set of methods are global methods that model all treatment effects and all possible inconsistency factors simultaneously, resulting in an omnibus test of incoherence in the whole network. The design-by-treatment interaction test is such a global method for incoherence (41,42). An overview of other methods for testing incoherence can be found elsewhere (40,48).

### The CINeMA approach

Both global and local incoherence tests have low power (49,50) and it is therefore important to consider the inconsistency factors as well as their uncertainty. As a large inconsistency factor may be indicative of a biased direct or indirect estimate, judging its magnitude is always important. As for imprecision and heterogeneity, the CINeMA approach to incoherence considers the impact on clinical conclusions, based on visual inspection of the 95% confidence interval of direct and indirect odds ratios and the range of equivalence. Consider the hypothetical examples in Figure 3 (Incoherence). The inconsistency factor using the SIDE splitting approach is the same for the three examples (1.27 with confidence interval 1.05 to 1.55), but their position relative to the range of equivalence differs and affects the interpretation of incoherence. In the first example, the 95% confidence intervals of both direct and indirect odds ratios lie above the range of equivalence: treatment A is clearly favourable, and there are ‘no concerns’ regarding inconsistency. In the second example, the 95% confidence interval of the indirect odds ratio straddles the range of equivalence while for the direct odds ratio the 95% confidence interval lies entirely above 1.05. In this situation, a judgement of ‘some concerns’ is appropriate. In the third example, the odds ratios from direct and indirect comparisons are in opposite directions and the disagreement will therefore lead to an expression of ‘major concerns’.

Note that in the three hypothetical examples above, both direct and indirect estimates exist. It could be, however, that there is only direct (e.g. venlafaxine versus vortioxetine in the network of antidepressants) or only indirect (e.g. agomelative versus vortioxetine) evidence. In this situation, we can neither estimate an inconsistency factor nor judge potential implications with respect to the range of equivalence. Considerations of indirectness and intransitivity are nevertheless important. Statistically, incoherence can only be judged using the global design-by-treatment interaction test. When a comparison is informed only by direct evidence, no disagreement between sources of evidence occurs and thus ‘no concerns’ for incoherence apply. If only indirect evidence is present then there will always be ‘some concerns’. There will be ‘major concerns’ if the p-value of the design-by-treatment interaction test is <0.01. As in comparisons informed only by indirect evidence coherence cannot be tested, having ‘no concerns’ for the particular treatment effects would be difficult to defend.

### Example: comparing antidepressants

In the network of antidepressants, the direct odds ratio comparing clomipramine with fluvoxamine is almost double the indirect odds ratio: the ratio of the two odds ratios (i.e., the inconsistency factor) is 1.94 (95% confidence interval 0.65 to 5.73, Table 4). However, both direct and indirect estimates contain values that extend to clinically important values in both directions. Thus, incoherence will not affect the interpretation of the NMA treatment effect: there are ‘no concerns’ (Table 5). In contrast, there are ‘major concerns’ regarding the confidence in the citalopram versus venlafaxine comparison: the direct odds ratio contains values within and above the range of equivalence while the indirect odds ratio includes values within and below the range of equivalence. The resulting estimated ratio of odds ratios is 2.08 (95% confidence interval 1.03 to 4.18) and the respective p-value of the SIDE test is 0.04 (Table 4). For the comparisons of amitriptyline versus paroxetine, the ratio of direct to indirect odds ratios is 1.05 (with 95% confidence interval (0.76, 1.46) and p-value 0.75) implying that the two sources of evidence are in agreement (Table 4). Direct and indirect estimates are very close in terms of odds ratios, 95% confidence intervals and the range of equivalence and we therefore have ‘no concerns’ regarding incoherence for this particular comparison.

## Summarizing judgments across the six domains

The output of the CINeMA framework is a table with the level of concern for each of the six domains. Some of the domains are interconnected: factors that may reduce the confidence in a treatment effect may affect more than one domain. Indirectness includes considerations on intransitivity, which manifest itself in the data as statistical incoherence. Heterogeneity may be related to most of the other domains. Pronounced heterogeneity will increase imprecision in treatment effects and may be related to variability in within-study biases or the presence of publication bias. Finally, in the presence of heterogeneity the ability to detect important incoherence will decrease (49).

Although the final output of CINeMA is a table with the level of concern for each of the six domains, reviewers may choose to summarize judgements across domains. If such an overall assessment is required, one may use the four levels of confidence using the usual GRADE approach: ‘very low’, ‘low’, ‘moderate’ or ‘high’ (24). For this purpose, an initial strategy would be to start at ‘high’ confidence and to drop a rating for each domain with some concerns and to drop two levels for each domain with major concerns. However, the six CINeMA domains should be considered jointly rather than in isolation, avoiding downgrading the overall level of confidence more than once for related concerns. For example, for the ‘citalopram versus venlafaxine’ comparison, we have ‘some concerns’ for imprecision and heterogeneity and ‘major concerns’ for incoherence (Table 3). However, downgrading by two levels will be sufficient in this situation, because imprecision, heterogeneity and incoherence are interconnected.

## Discussion

We have outlined and illustrated the CINeMA approach for evaluating confidence in treatment effect estimates from NMA, covering the six domains of within-study bias, across-study bias, indirectness, imprecision, heterogeneity and incoherence. Our approach avoids selective use of indirect evidence, while considering the characteristics of all studies included in the network. Thus, we are not using assessments of confidence to decide whether to present direct or indirect (or combined) evidence, as has been suggested by others (4,5). We differentiate between the three sources of variability in a network, namely, imprecision, heterogeneity and incoherence and we consider the impact that each source might have on decisions for treatment. The approach can be operationalized and is easy-to-implement even for very large networks.

Any approach to evaluating confidence in evidence synthesis results will inevitably involve some subjectivity. Our approach is no exception. While the use of bar charts to infer about the impact of within study biases and indirectness provides a consistent assessment across all comparisons in the network, their summary is difficult. Setting up a margin of equivalence might be equivocal. Further limitations of the framework are associated with the fact that published articles are used to make judgements and these reports do not necessarily reflect the way studies were undertaken. For instance, judging indirectness requires study data to be collected on pre-specified effect modifiers; reporting limitations will inevitably impact on the reliability of the judgements.

A consequence of the inherent subjectivity of the system is that interrater agreement may be modest. Studies of the reproducibility of assessments made by researchers using CINeMA will be required in this context. We believe however that transparency is key. Although judgements may differ across reviewers, they are made using explicit criteria. These should be specified in the review protocol so that data-driven decisions are avoided. The web application at *cinema.ispm.ch* will greatly facilitate the implementation of all steps involved in the application of CINeMA (6).

This paper proposes a refinement of a previously suggested framework (51). An alternative approach has also been refined (52) since its initial introduction (53). The two methods have similarities but also notable differences. For example, Puhan et al (53) suggest a process of deciding whether indirect estimates are of sufficient certainty to combine them with the direct estimates. In contrast CINeMA evaluates relative treatment effects without considering separately the direct and indirect sources. Evaluation of the impact of within-study bias and indirectness differs materially between the two approaches. The need to choose the most influential one-step loop in the GRADE approach as described by Puhan et al. (53) and Brignardello-Petersen (18) discards a large amount of information that contributes to the results and makes the approach difficult to apply to large networks. The percentage contribution matrix appears to be the only viable option to acknowledge the impact of each and every study included in a network. Moreover, our framework naturally includes the results from sensitivity analyses in the interpretation of the bar charts. Finally, in contrast to GRADE approach, we do not rely on metrics for judging heterogeneity and incoherence: we consider instead the impact that these can have when a stakeholder needs to make informed decisions. An alternative approach to assessing confidence findings from network meta-analysis is to explore how robust treatment recommendations are to potential degrees of bias in the evidence (54). The method is easy to apply but focuses on the impact of bias and does not explicitly address heterogeneity, indirectness and inconsistency.

Evidence synthesis is increasingly used by national and international agencies (55,56) to inform decisions about the reimbursement of medical interventions, by clinical guideline panels to recommend one drug over another and by clinicians to prescribe a treatment or recommend a diagnostic procedure for individual patients. However, it is the exception rather than the rule for published network meta-analyses to formally evaluate confidence in relative treatment effects (57). With the use of open-source free software (see Box 2), our approach can be routinely applied to any network meta-analysis (6) and offers a step forward in transparency and reproducibility. The suggested framework operationalizes, simplifies and accelerates the process of evaluation of results from large and complex networks without compromising in statistical and methodological rigor. The CINeMA framework is a transparent, rigorous and comprehensive system for evaluating the confidence of treatment effect estimates from network meta-analysis.

### Box 2.

Description of the CINeMA web-application.

#### THE CINeMA WEB APPLICATION

CINeMA framework has been implemented in a freely available, user-friendly web-application aiming to facilitate the evaluation of confidence on the results from network meta-analysis (http://cinema.ispm.ch/ (6)). The web application is programmed in javascript, uses docker and is linked with R; in particular, packages *meta* and *netmeta* are used (59). Knowledge of the aforementioned languages and technologies is however not required from the users.

##### Loading the data

In ‘My Projects’ tab, CINeMA users are able to upload a .csv file with the by-treatment outcome study data and study-level risk of bias (RoB) and indirectness judgments. CINeMA web-application can handle all the formats used in network meta-analysis (long or wide format, binary or continuous, arm level or study level data) and provides flexibility in labelling variables as desired by the user. A demo dataset is available in ‘My Projects’ tab.

##### Evaluating the confidence in the results from network meta-analysis

A preview of the evidence (network plot and outcome data) and options concerning the analysis (fixed or random effects, effect measure etc.) are given in the ‘Configuration’ tab. The next six tabs guide users to make informed conclusions on the quality of evidence based on within-study bias, across-studies bias, indirectness, imprecision, heterogeneity and incoherence. Features implemented include the percentage contribution matrix, relative treatment effects for each comparison, estimation of the heterogeneity variance, prediction intervals and tests for the evaluation of the assumption of coherence.

##### Summarising judgments

The last tab ‘Report’ includes a summary of the evaluations made in the six domains and gives users the possibility to either not downgrade, or downgrade by one or two levels each relative treatment effect. Users can download a report with the summary of their evaluations along with their final judgements. CINeMA is accompanied by a documentation describing each step in detail (tab ‘Documentation’).

## Acknowledgements

The development of the software and part of the presented work was supported by the Campbell Collaboration. ME was supported by special project funding (Grant No. 174281) from the Swiss National Science Foundation. GS, AN, TP were supported by project funding (Grant No. 179158) from the Swiss National Science Foundation. The funders had no role in study design, data collection and analysis, decision to publish, or preparation of the manuscript.

## Supplementary Material

**Table S1.**
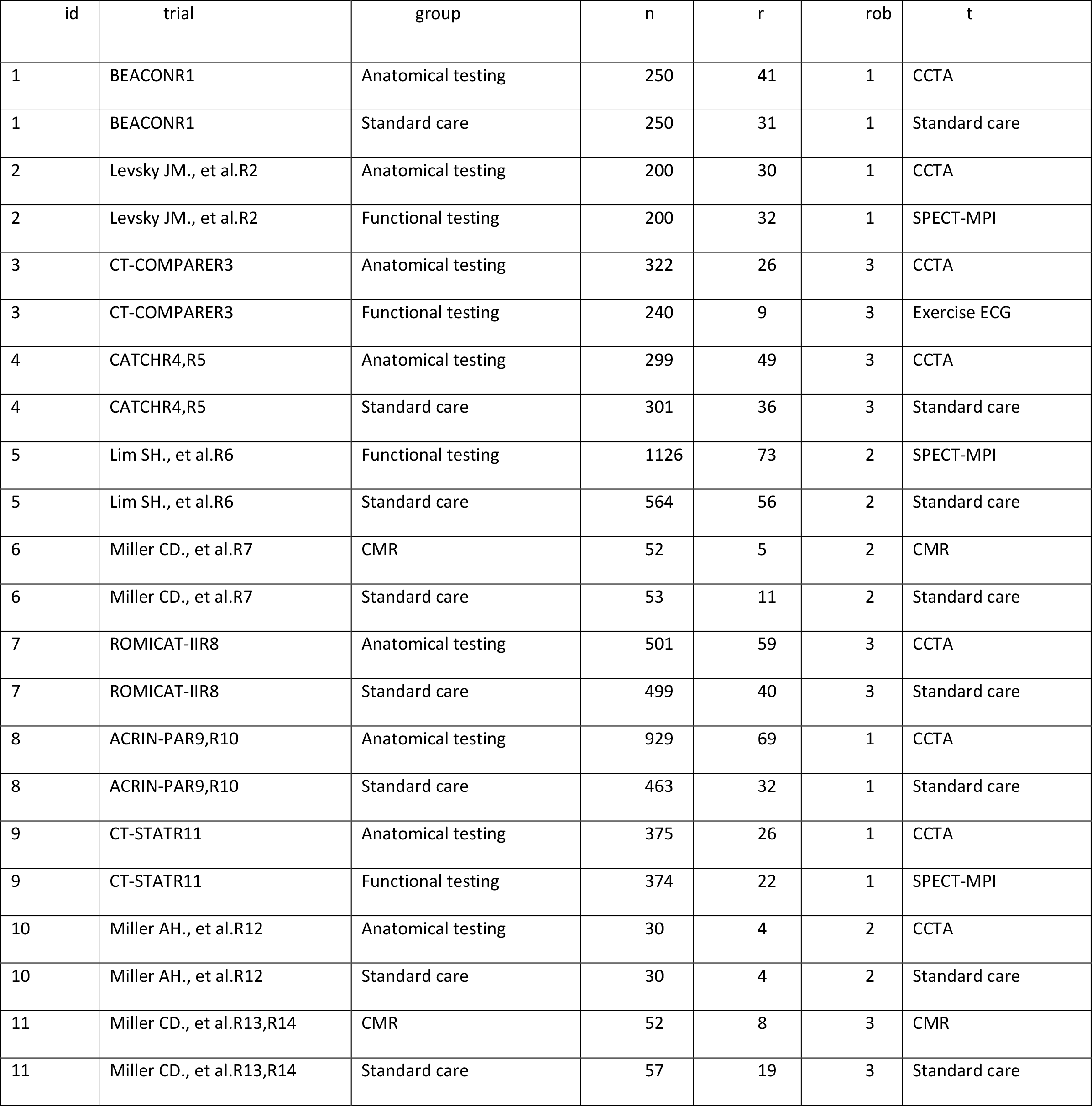
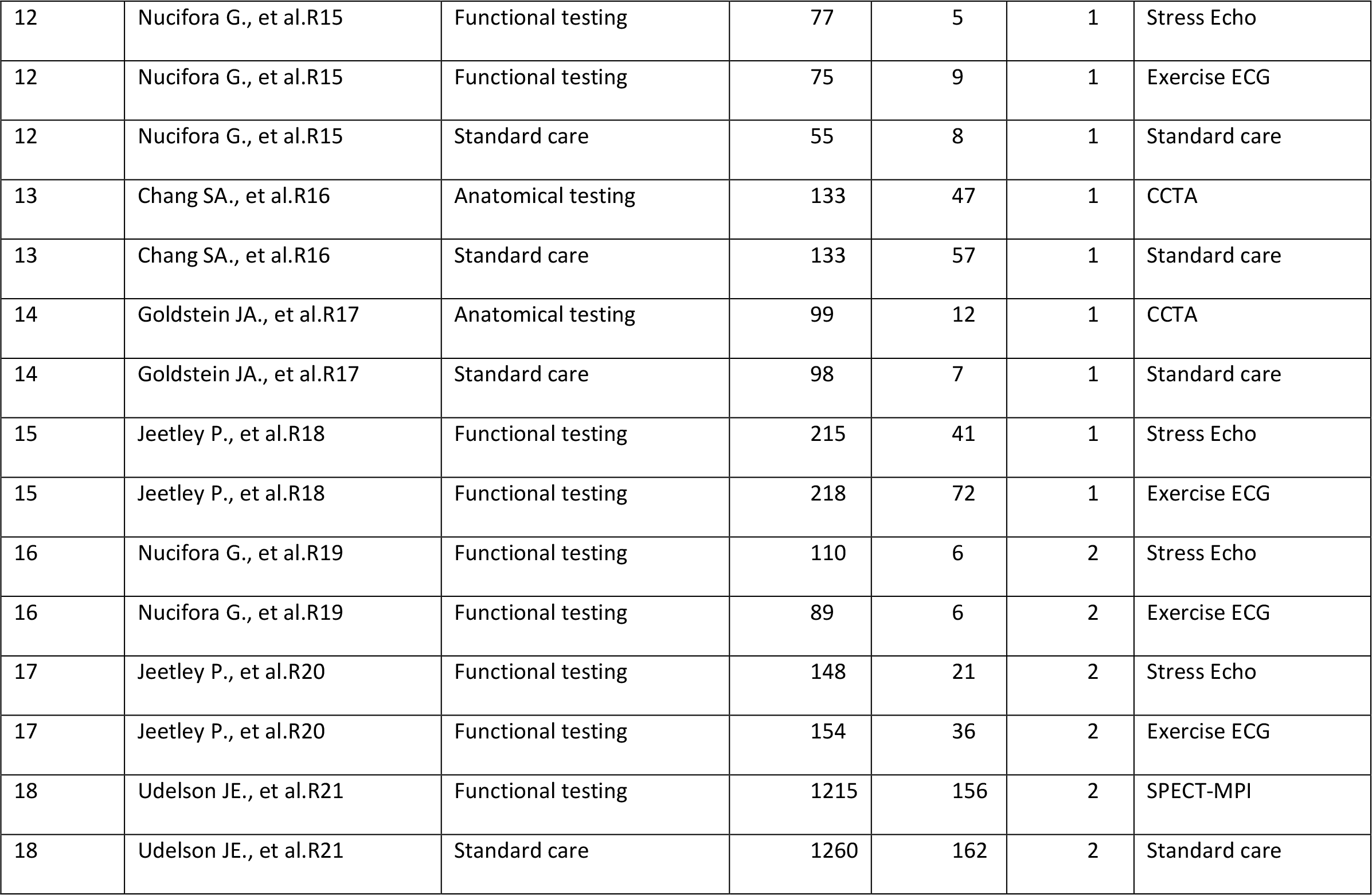
Data from Network of randomised controlled trials comparing non-invasive diagnostic strategies for the detection of coronary artery disease in patients with low risk acute coronary syndrome. The data was originally published by Siontis et al.

| id | trial | group | n | r | rob | t |
| --- | --- | --- | --- | --- | --- | --- |
| 1 | BEACONR1 | Anatomical testing | 250 | 41 | 1 | CCTA |
| 1 | BEACONR1 | Standard care | 250 | 31 | 1 | Standard care |
| 2 | Levsky JM., et al.R2 | Anatomical testing | 200 | 30 | 1 | CCTA |
| 2 | Levsky JM., et al.R2 | Functional testing | 200 | 32 | 1 | SPECT-MPI |
| 3 | CT-COMPARER3 | Anatomical testing | 322 | 26 | 3 | CCTA |
| 3 | CT-COMPARER3 | Functional testing | 240 | 9 | 3 | Exercise ECG |
| 4 | CATCHR4,R5 | Anatomical testing | 299 | 49 | 3 | CCTA |
| 4 | CATCHR4,R5 | Standard care | 301 | 36 | 3 | Standard care |
| 5 | Lim SH., et al.R6 | Functional testing | 1126 | 73 | 2 | SPECT-MPI |
| 5 | Lim SH., et al.R6 | Standard care | 564 | 56 | 2 | Standard care |
| 6 | Miller CD., et al.R7 | CMR | 52 | 5 | 2 | CMR |
| 6 | Miller CD., et al.R7 | Standard care | 53 | 11 | 2 | Standard care |
| 7 | ROMICAT-IIR8 | Anatomical testing | 501 | 59 | 3 | CCTA |
| 7 | ROMICAT-IIR8 | Standard care | 499 | 40 | 3 | Standard care |
| 8 | ACRIN-PAR9,R10 | Anatomical testing | 929 | 69 | 1 | CCTA |
| 8 | ACRIN-PAR9,R10 | Standard care | 463 | 32 | 1 | Standard care |
| 9 | CT-STATR11 | Anatomical testing | 375 | 26 | 1 | CCTA |
| 9 | CT-STATR11 | Functional testing | 374 | 22 | 1 | SPECT-MPI |
| 10 | Miller AH., et al.R12 | Anatomical testing | 30 | 4 | 2 | CCTA |
| 10 | Miller AH., et al.R12 | Standard care | 30 | 4 | 2 | Standard care |
| 11 | Miller CD., et al.R13,R14 | CMR | 52 | 8 | 3 | CMR |
| 11 | Miller CD., et al.R13,R14 | Standard care | 57 | 19 | 3 | Standard care |
| 12 | Nucifora G., et al.R15 | Functional testing | 77 | 5 | 1 | Stress Echo |
| 12 | Nucifora G., et al.R15 | Functional testing | 75 | 9 | 1 | Exercise ECG |
| 12 | Nucifora G., et al.R15 | Standard care | 55 | 8 | 1 | Standard care |
| 13 | Chang SA., et al.R16 | Anatomical testing | 133 | 47 | 1 | CCTA |
| 13 | Chang SA., et al.R16 | Standard care | 133 | 57 | 1 | Standard care |
| 14 | Goldstein JA., et al.R17 | Anatomical testing | 99 | 12 | 1 | CCTA |
| 14 | Goldstein JA., et al.R17 | Standard care | 98 | 7 | 1 | Standard care |
| 15 | Jeetley P., et al.R18 | Functional testing | 215 | 41 | 1 | Stress Echo |
| 15 | Jeetley P., et al.R18 | Functional testing | 218 | 72 | 1 | Exercise ECG |
| 16 | Nucifora G., et al.R19 | Functional testing | 110 | 6 | 2 | Stress Echo |
| 16 | Nucifora G., et al.R19 | Functional testing | 89 | 6 | 2 | Exercise ECG |
| 17 | Jeetley P., et al.R20 | Functional testing | 148 | 21 | 2 | Stress Echo |
| 17 | Jeetley P., et al.R20 | Functional testing | 154 | 36 | 2 | Exercise ECG |
| 18 | Udelson JE., et al.R21 | Functional testing | 1215 | 156 | 2 | SPECT-MPI |
| 18 | Udelson JE., et al.R21 | Standard care | 1260 | 162 | 2 | Standard care |

**Table S2.**
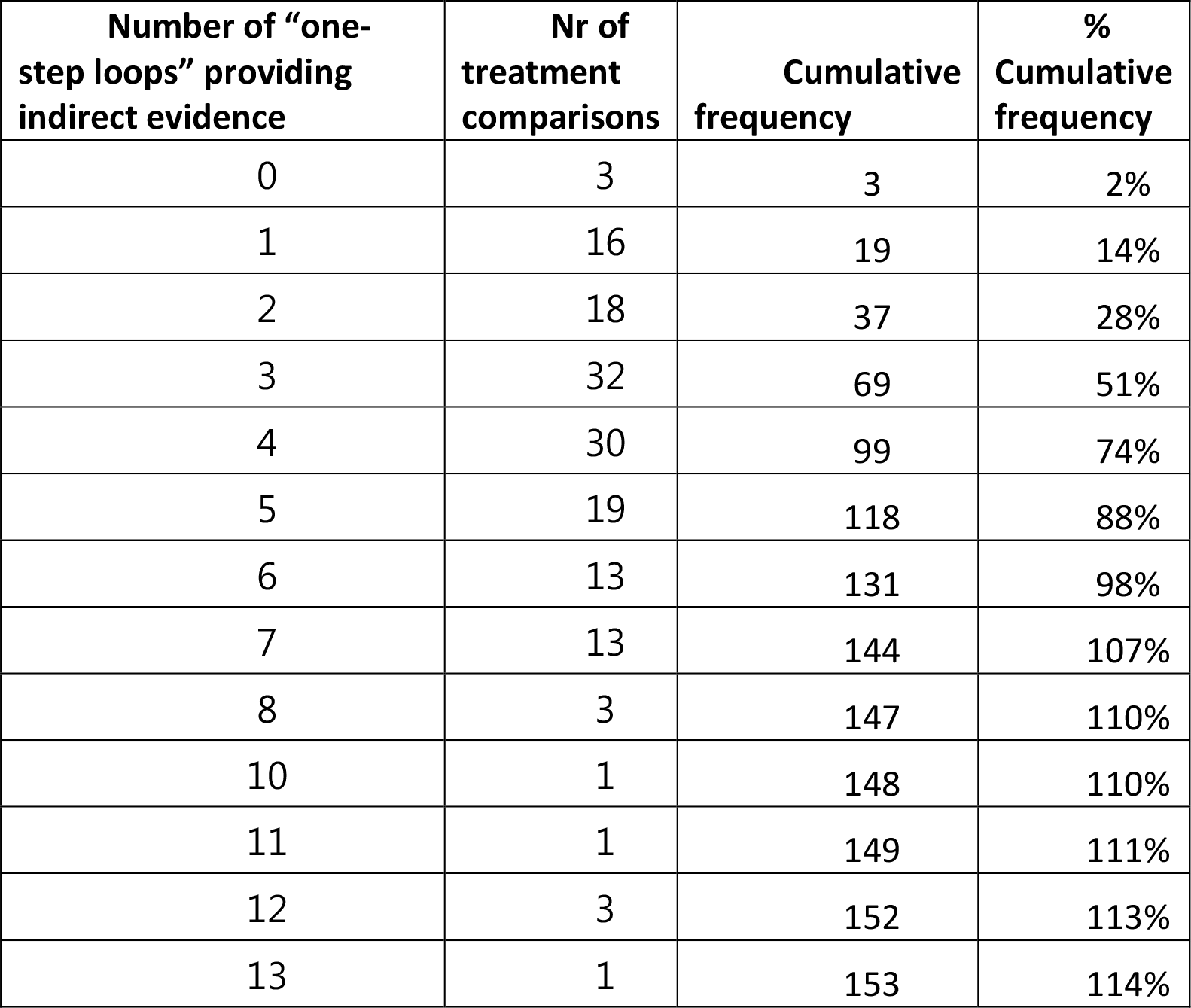
Number of "one-step loops" providing indirect evidence for NMA relative treatment effects between treatment comparisons for the network of antidepressants.

**Table S3.** Average contribution to NMA relative treatment effects from direct evidence and indirect evidence via intermediate comparators (steps). The “one-step loop” provides one-step indirect comparison via a single common treatment.

| Source of evidence |  | % Contribution |
| --- | --- | --- |
| Direct evidence |  | 11.1% |
| Indirect evidence | 1 step | 56.1% |
|  | 2 steps | 84.3% |
|  | 3 steps | 95.0% |
|  | 4 steps | 98.0% |
|  | 5 steps | 99.1% |
|  | 6 steps | 99.5% |
|  | 7 steps | 99.7% |
|  | 8 steps | 99.9% |
|  | 9 steps | 100.0% |
|  | 10 steps | 100.0% |
|  | 11 steps | 100.0% |
|  | 12 steps | 100.0% |
|  | 13 steps | 100.0% |

**Table S4.**
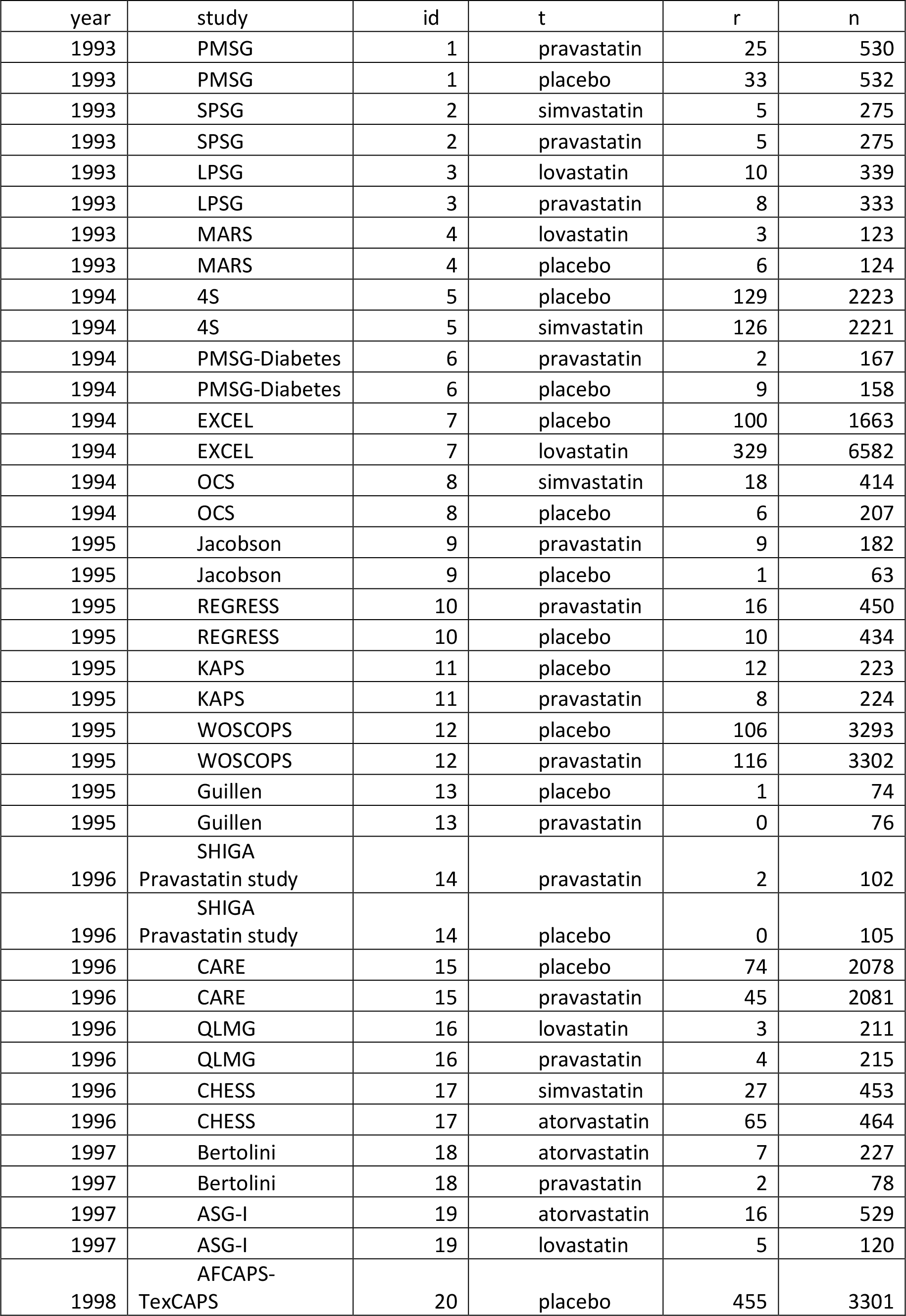
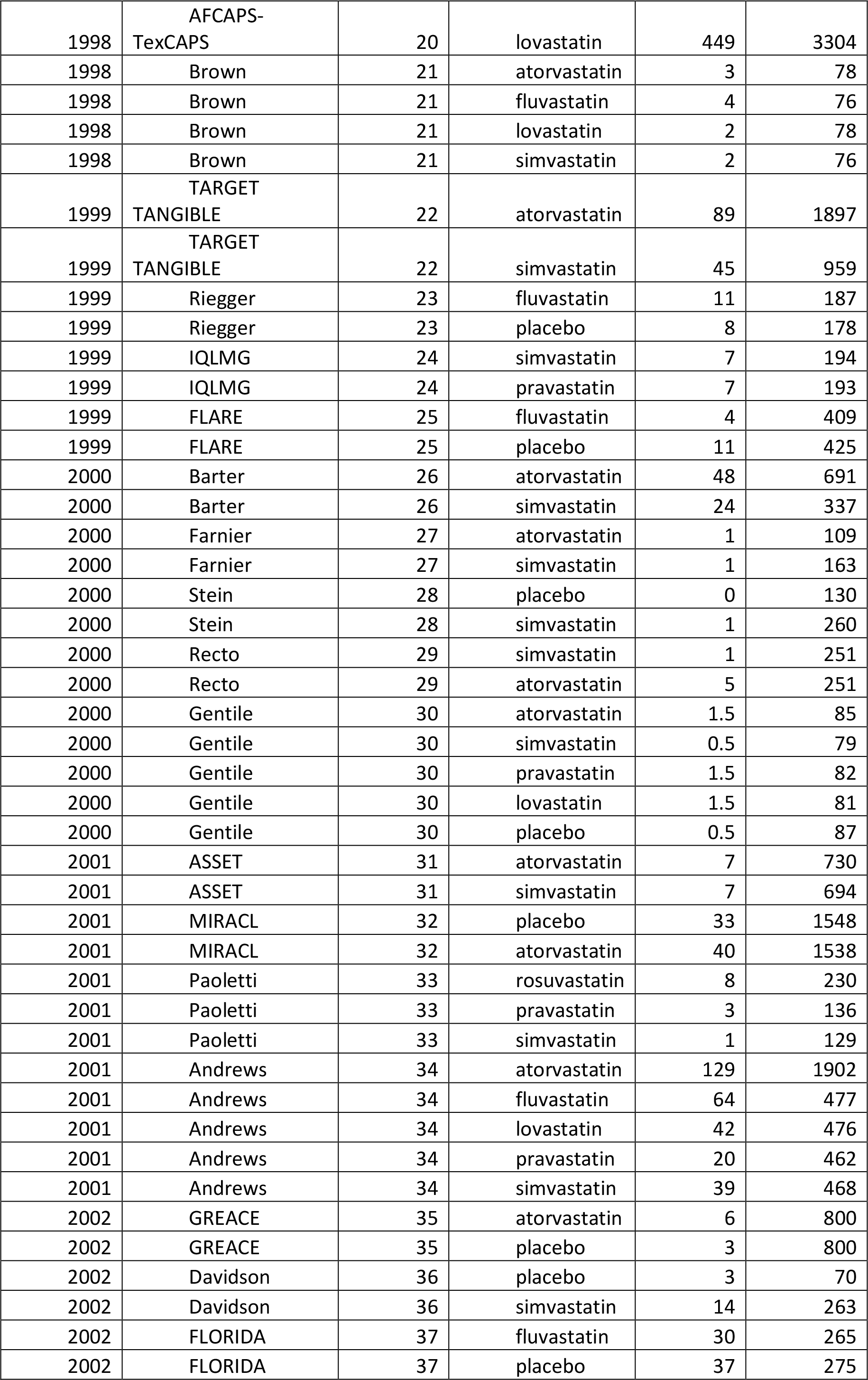
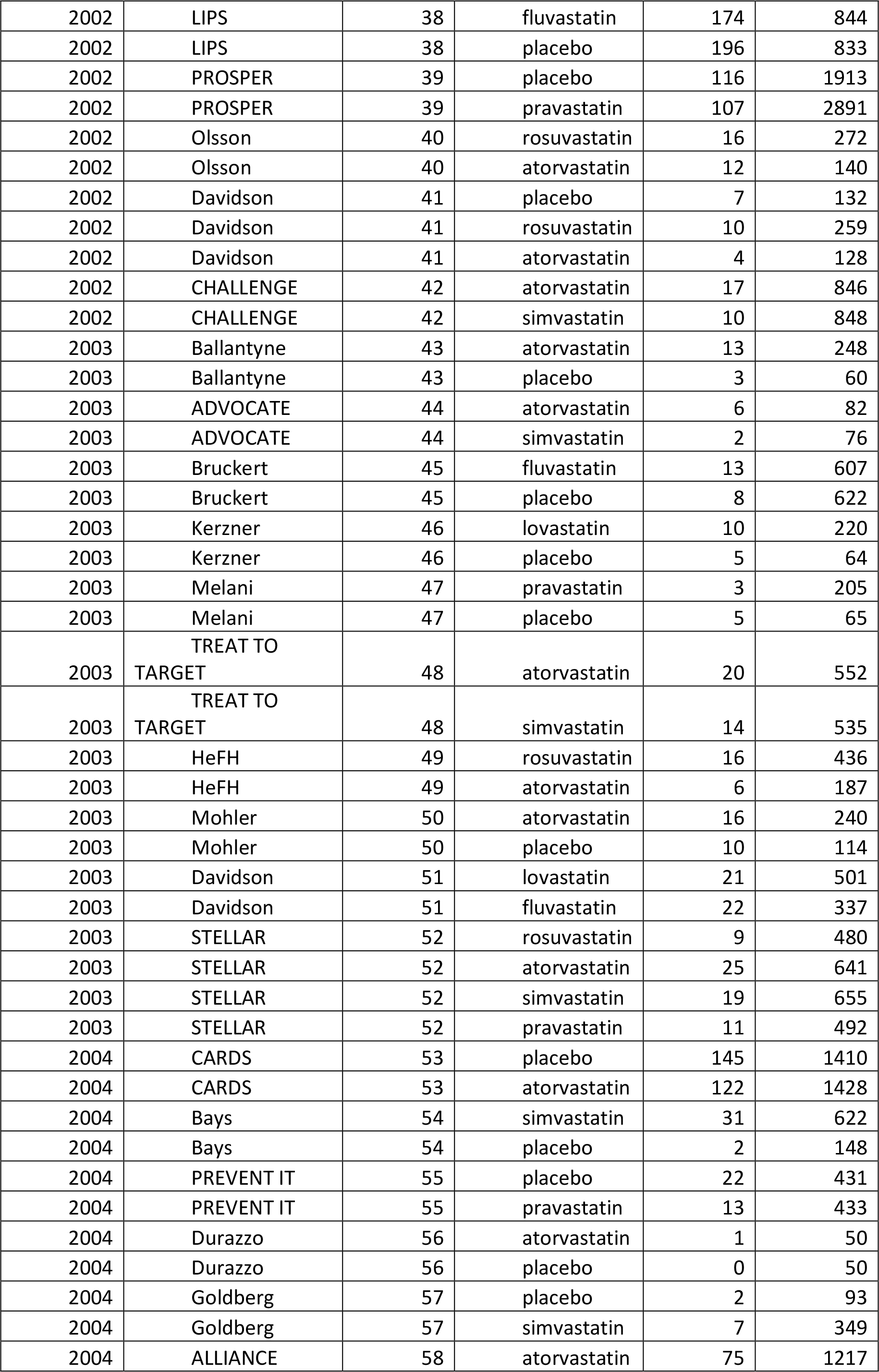
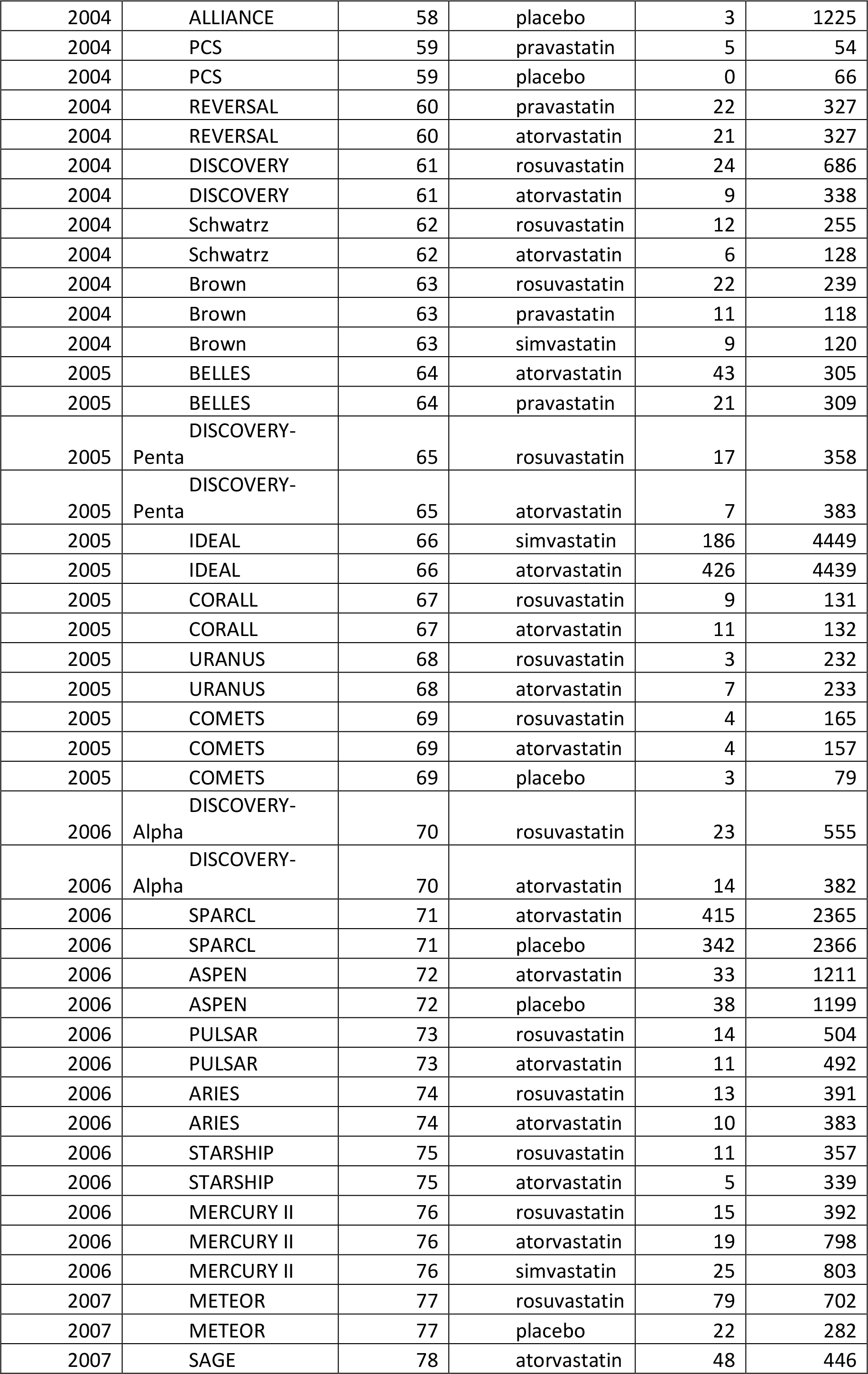
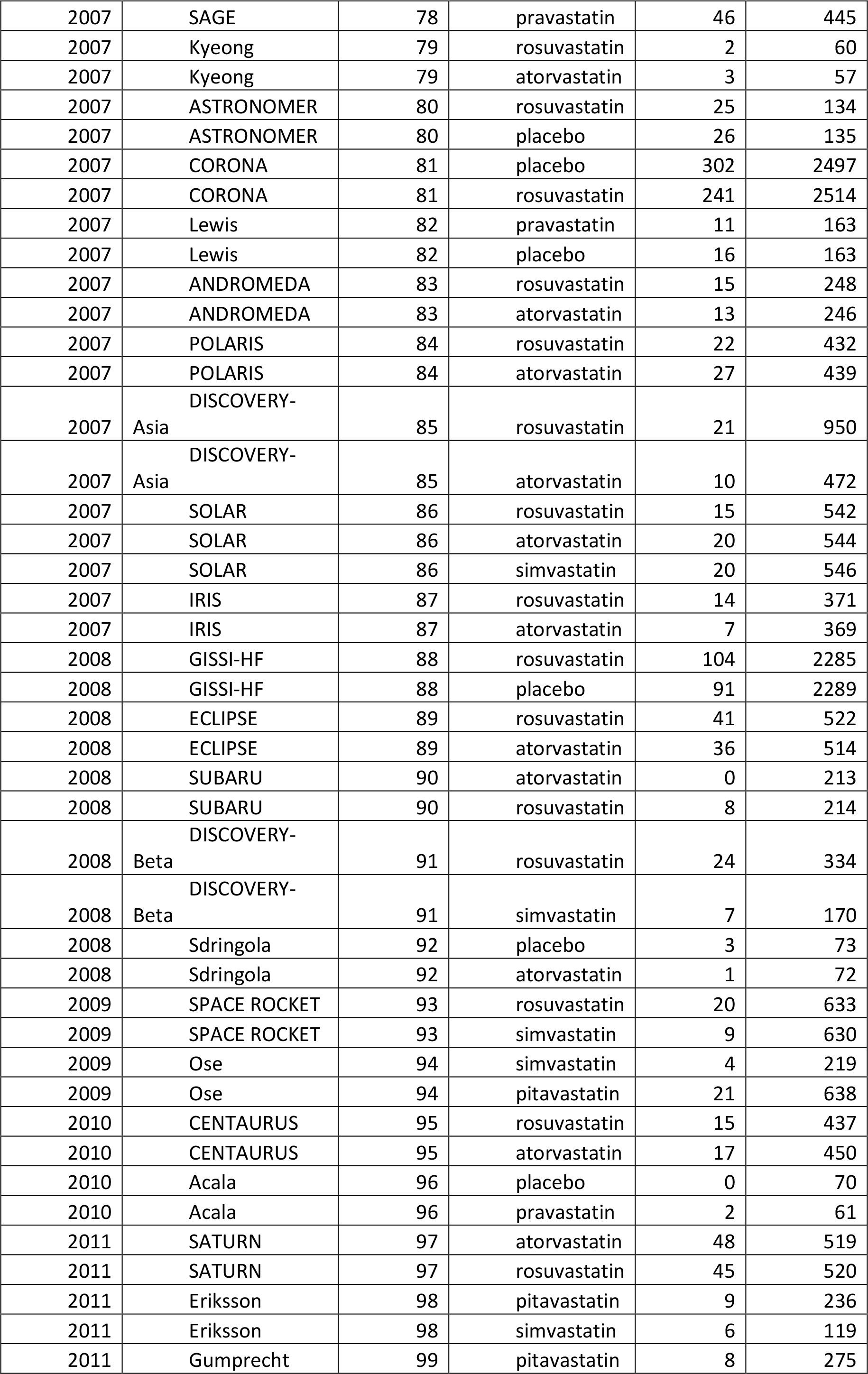
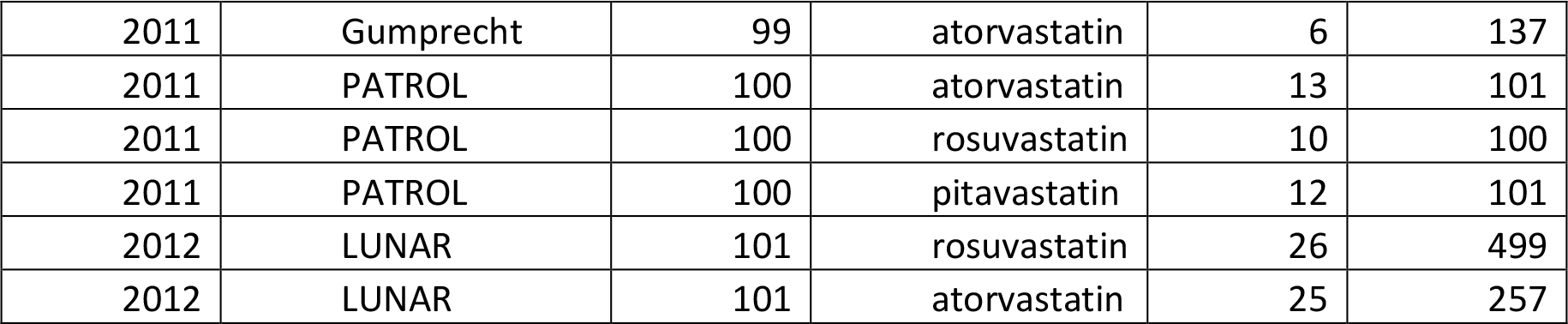
Data from the network of randomised controlled trials comparing adverse effects of statins. The data was originally published by Naci et al. id: id of the study, t: treatment name, r: number of adverse effects, n: sample size.

**Table S5.** NMA results from the network of randomised controlled trials comparing adverse effects of statins. Odds ratios and their 95% confidence intervals are presented. Odds ratios less than 1 favor the treatment specified in the row.

|  |  |  |  |  |  |  |  |
| --- | --- | --- | --- | --- | --- | --- | --- |
| **Atorvastatin** | 0.894<br>(0.637, 1.255) | 1.196<br>(0.868, 1.647) | 1.127<br>(0.637, 1.994) | 1.073<br>(0.890, 1.294) | 1.418<br>(1.126, 1.785) | 1.007<br>(0.848, 1.196) | 1.297<br>(1.065, 1.580) |
| 1.119<br>(0.797, 1.571) | **Fluvastatin** | 1.338<br>(0.915, 1.956) | 1.261<br>(0.655, 2.426) | 1.201<br>(0.879, 1.640) | 1.586<br>(1.109, 2.268) | 1.127<br>(0.787, 1.612) | 1.451<br>(1.011, 2.083) |
| 0.836<br>(0.607, 1.152) | 0.747<br>(0.511, 1.093) | **Lovastatin** | 0.942<br>(0.494, 1.797) | 0.897<br>(0.668, 1.206) | 1.185<br>(0.849, 1.655) | 0.842<br>(0.599, 1.184) | 1.085<br>(0.768, 1.532) |
| 0.887<br>(0.501, 1.570) | 0.793<br>(0.412, 1.526) | 1.061<br>(0.557, 2.023) | **Pitavastatin** | 0.952<br>(0.528, 1.718) | 1.258<br>(0.687, 2.305) | 0.893<br>(0.499, 1.598) | 1.151<br>(0.648, 2.043) |
| 0.932<br>(0.773, 1.123) | 0.833<br>(0.610, 1.138) | 1.114<br>(0.829, 1.498) | 1.050<br>(0.582, 1.895) | **Placebo** | 1.321<br>(1.070, 1.632) | 0.938<br>(0.759, 1.160) | 1.209<br>(0.960, 1.522) |
| 0.705<br>(0.560, 0.888) | 0.630<br>(0.441, 0.902) | 0.844<br>(0.604, 1.178) | 0.795 (0.434,<br>1.457) | 0.757<br>(0.613, 0.935) | **Pravastatin** | 0.710<br>(0.549, 0.918) | 0.915<br>(0.702, 1.193) |
| 0.993<br>(0.836, 1.179) | 0.888<br>(0.620, 1.270) | 1.188<br>(0.844, 1.671) | 1.119<br>(0.626, 2.002) | 1.066<br>(0.862, 1.317) | 1.408<br>(1.089, 1.821) | **Rosuvastatin** | 1.288<br>(1.024, 1.621) |
| 0.771<br>(0.633, 0.939) | 0.689<br>(0.480, 0.989) | 0.922<br>(0.653, 1.302) | 0.869<br>(0.489, 1.542) | 0.827<br>(0.657, 1.041) | 1.093<br>(0.839, 1.424) | 0.776<br>(0.617, 0.977) | **Simvastatin** |

**Table S6.**
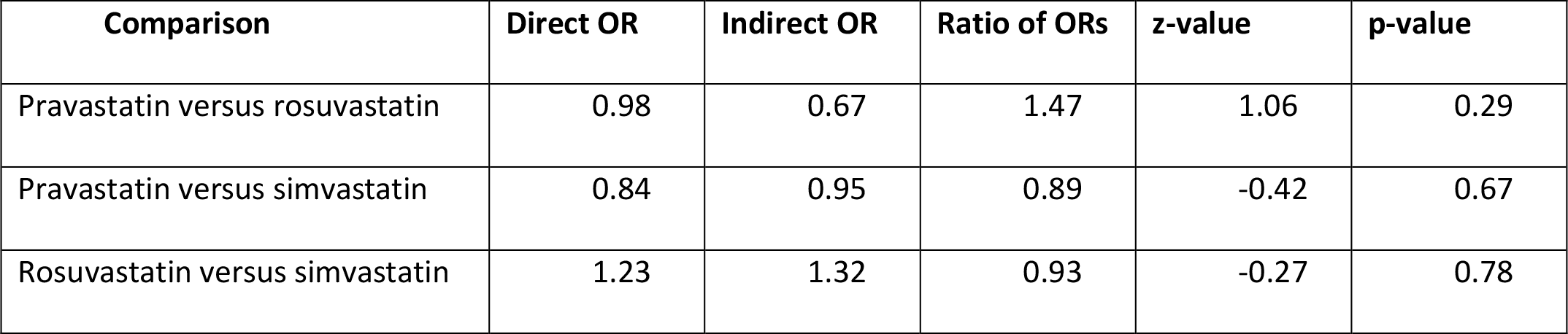
Results from SIDE splitting for three network comparisons of the network of statins. OR: odds ratio. SIDE: separate indirect from direct approach.

| Comparison | Direct OR | Indirect OR | Ratio of ORs | z-value | p-value |
| --- | --- | --- | --- | --- | --- |
| Pravastatin versus rosuvastatin | 0.98 | 0.67 | 1.47 | 1.06 | 0.29 |
| Pravastatin versus simvastatin | 0.84 | 0.95 | 0.89 | -0.42 | 0.67 |
| Rosuvastatin versus simvastatin | 1.23 | 1.32 | 0.93 | -0.27 | 0.78 |

